# The curse of zombie dispersal in discrete-time models of spatial population dynamics

**DOI:** 10.1101/2023.10.31.565055

**Authors:** Michael G. Neubert, Silke F. van Daalen

## Abstract

In many metapopulation and metacommunity models, individuals disperse between discrete habitat patches. When those models treat time as a discrete variable, the formulation of the dispersal term must be handled with care. A commonly made mistake is to model dispersal with terms identical to those found in continuous-time models. Such terms can inadvertently resurrect dead individuals, effectively creating “zombie dispersers.” Zombie dispersal, in turn, can have dramatic, but spurious, effects on model dynamics. In this manuscript, we illustrate the misleading effects generated by zombie dispersal in a published model used to investigate how dispersal mediates synchrony in population dynamics.

## Introduction

Dispersal—the movement of individuals in space—along with reproduction and survival, is one of the fundamental processes that determine population dynamics. Dispersal affects how much habitat is required for a population to persist, the rate at which a population will spread when introduced into new habitat, and the degree to which populations separated in space exhibit synchronous fluctuations. In metapopulations, local populations are confined to habitat patches and are subject to extinction (Hanski, 1999) and colonization from populations in other habitat patches. The rate of colonization is determined by dispersal, and colonization rates determine whether the metapopulation will persist.

In mathematical models of metapopulation dynamics, the description of dispersal depends on the the description of time. Consider a simple model for a metapopulation distributed in two patches: one labeled *x*, the other *y*. If time (*t*) is treated as a continuous variable, and as a result the population sizes in the two patches (*N*_*x*_(*t*) and *N*_*y*_(*t*)) are described using differential equations, the model would take the form

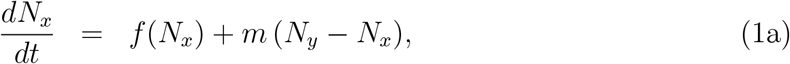

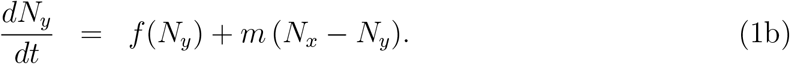

In this model, the function *f* describes the dependence of survival and reproduction on local population size and *m* is the (non-negative) emigration rate—the rate at which individuals leave one patch and arrive at the other.

In other models, time is treated as a discrete variable, taking on only integer values. Discrete-time population models are common in mathematical demography (Keyfitz and Caswell, 2005) for populations living in strongly seasonal environments in which births and deaths are typically concentrated in time (Kot, 2001). In the physics and mathematics literature, discrete-time, discrete-space models are known as “coupled map lattices” (CML) (Kaneko, 1993).

It is tempting to formulate dispersal in a CML metapopulation model in analogy with its continuous-time counterpart (1); that is, as^1^

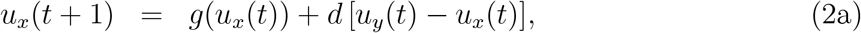

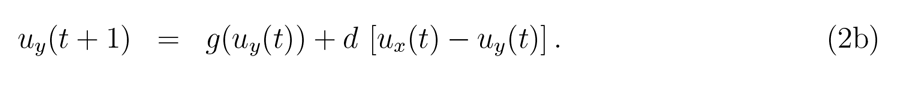

CMLs with this kind of interpatch coupling have been used to study the synchronization of coupled oscillators (Poria et al., 2008; Poria et al., 2013; Mehta and Sinha, 2000; Khan and Sahoo, 2016; Nag and Poria, 2020; Khan et al., 2022). They have also been used in ecology to study spatial population dynamics (Vance, 1984; Bascompte and Solé, 1994) including the synchronization of population fluctuations across spatially distinct habitat patches (Shoemaker et al., 2022).

The formulation of dispersal in model (2), however, is biologically problematic. In this brief paper we review how this **zombie dispersal** problem arises and how it can lead to incorrect conclusions about the effects of metapopulation and metacommunity dynamics.

It is easiest to illustrate the problem with the formulation of dispersal in model (2) by presenting a special case where

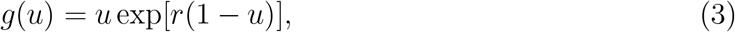

and when the population is initially concentrated in patch *x* (i.e., *𝓊*_*y*_(0) = 0). In this case, *𝓊*_*x*_(1) will be negative if 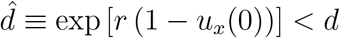, a condition that is likely to occur when *r* is large (so intraspecific competition is intense) and dispersal is high.

The possibility of negative population size arises because model (2) violates a property of logically-consistent discrete-time models: only one process should occur at a time. In model (2), some individuals die, but are then counted again in the adjoining population as the result of an unrealistic, zombie dispersal process.^2^

Even when 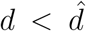, the dynamics of a model with zombie dispersal, like (2), can generate misleading results, especially when coupled with strong nonlinearities (e.g., over-compensation or Allee effects). We can see this by comparing the dynamics of (2) with a model that does not include the undead among the dispersers.

One correct way to model dispersal is to specify that a certain fraction (*d*) of the individuals *that remain after reproduction and mortality* emigrate to the adjoining patch, as in:

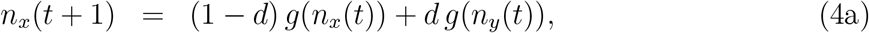

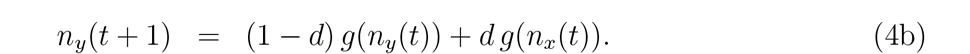

where *g* is again given by (3).^3^

Both models (2)-(3) and (4)-(3) have a spatially uniform equilibrium of the form *𝓊*_*x*_(*t*) = *𝓊*_*y*_(*t*) = *𝓊*^*\**^ = 1 and *n*_*x*_(*t*) = *n*_*y*_(*t*) = *n*^*\**^ = 1; however, the stability of these equilibria, and how it depends on model parameters, is very different in the two models. A simple analysis shows that *n*^*\**^ is stable if *r <* 2, and that *𝓊*^*\**^ is stable if *r <* 2(1 *− d*). That is, while dispersal does not affect stability, zombie dispersal is destabilizing.

In addition, when both equilibria are unstable, the asymptotic dynamics can be very different (Fig. 1). For example, when *r* = 2.1 and *d* = 0.2, the zombie dispersal model is asymptotically chaotic (Fig. 1A) and the trajectories of population density in the two patches are out of phase (Fig. 1B). The non-zombie model exhibits a two-cycle (Fig. 1C) with the two populations synchronized (Fig. 1D).

**Figure 1.**
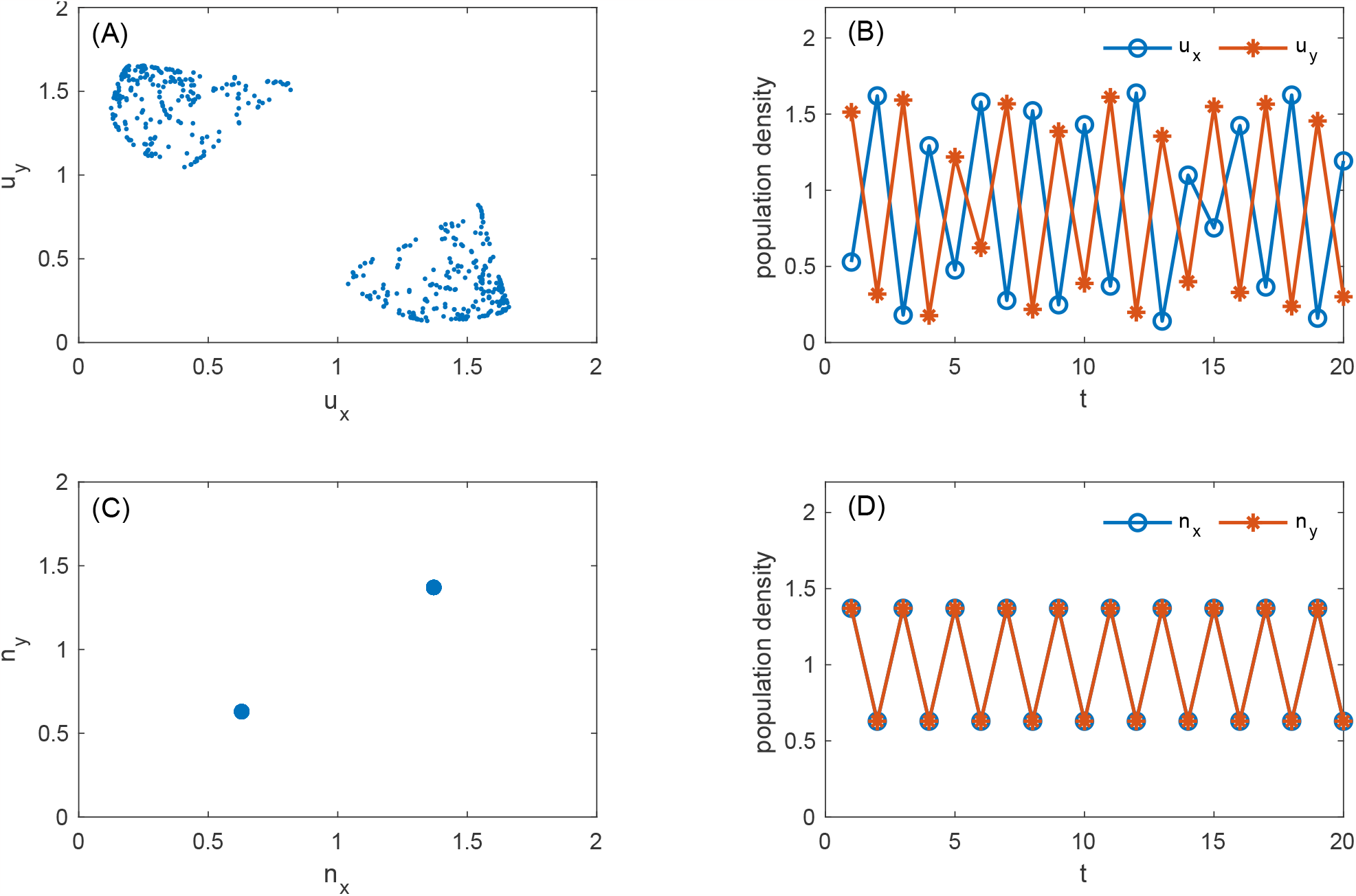
Simulations of models (2)-(3) in panels (A) and (B), and (4)-(3) in panels (C) and (D). *r* = 2.1 and *d* = 0.2 in both simulations. The asymptotic attractor of the zombie-dispersal model (2) is strange (A) and the trajectory is chaotic (B) with the two populations fluctuating out of phase. In contrast, the asymptotic attractor of model (4) is a two-cycle (C), and the two populations fluctuate synchronously (D).

The differences in the synchronizing effects of dispersal versus zombie dispersal can have important ecological implications (Earn et al., 2000; Earn and Levin, 2006). Synchrony in the dynamics of constituent populations typically reduces the persistence and resilience of the ecological communities and metacommunities they comprise (Heino et al., 1997; Micheli et al., 1999; Lamy et al., 2021). Compensatory community dynamics, in which the abundances of constituent populations are negatively correlated, tends to increase stability and reduce variability in aggregate quantities such as total community biomass.

### Zombie Dispersal and Metacommunity Synchrony

In a recent paper, Shoemaker et al. (2022) used a diagnostic index of synchrony to study how various ecological processes, including dispersal, affect synchrony on both short and long time scales. In this section and the next, we point out that the model of dispersal in Shoemaker et al. (2022) includes zombies, and show that this inconsistent model can lead to incorrect conclusions about the effects of dispersal on synchrony.

One diagnostic measure of synchrony between the time series of population abundances in a community (call them *n*_*i*_(*t*)) is the variance ratio

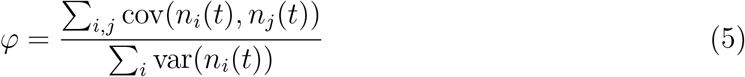

(Shluter, 1984). Synchrony is indicated when *φ >* 1; compensation when *φ <* 1. Zhao et al. (2020) pointed out that communities can exhibit synchrony (or compensation) at different timescales that cannot be distinguished by the classic variance ratio (5), but can be distinguished by the “timescale-specific variance ratio”

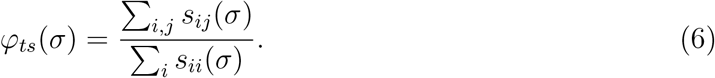

Here, *s*_*ii*_(*σ*) is the power spectral density of *n*_*i*_ and *s*_*ij*_(*σ*) is the cospectrum of *n*_*i*_(*t*) and *n*_*j*_(*t*). A value of *φ*_*ts*_(*σ*) larger than one indicates that the two populations exhibit synchronous dynamics at the timescale *σ*; *φ*_*ts*_(*σ*) *<* 1 indicates compensatory dynamics.

Shoemaker et al. (2022) applied the timescale-specific variance ratio (6) to the output of a family of mathematical models to understand how several ecological processes might affect synchrony on both short and long timescales. They also compared the timescale-specific results to the classic ratio (5). They found that “… synchronous and compensatory dynamics are not mutually exclusive but instead can vary by timescale” and that the classic ratio is “often biased toward long-term drivers and may miss the importance of short-term drivers.”

Dispersal was one of the processes that Shoemaker et al. (2022) examined, with the following model, which tracks the population densities of two competing species in two habitat patches. If the population density of species *i* in patch *x* at time *t* is denoted as *N*_*i,x*_(*t*), and similarly the population density of species *j* in patch *y* as *N*_*j,y*_(*t*), the model takes the form

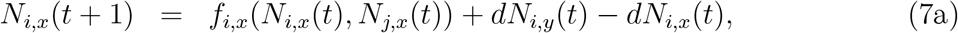

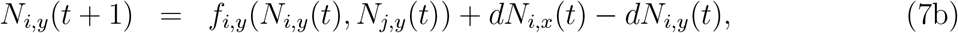

where

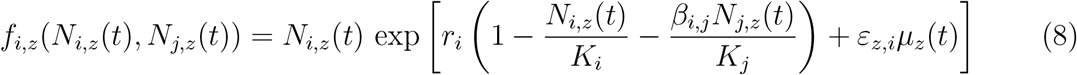

for *i* ∈ {1, 2}; *z* ∈ {*x, y*}; and *j* ≠ *i*.

In the absence of dispersal, when *d* = 0, the model assumes that the density in patch *z* changes as the result of survival and reproduction, at a natural per capita rate 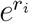. The rate is reduced by intraspecific density dependence (exp(*−r*_*i*_*N*_*i,z*_(*t*)*/K*_*i*_)) and interspecific competition (exp(*−r*_*i*_*β*_*i,j*_*N*_*j,z*_(*t*)*/K*_*j*_)). The local population growth rate of species *i* in patch *z* is also modified by periodic environmental fluctuations (exp(*ε*_*z,i*_*μ*_*z*_(*t*)), where *μ*_*z*_(*t*) = *a*_*z*_ sin(*b*_*z*_*t* + *c*_*z*_)). At the same time, the model posits that some fraction *d* of the individuals of both species in both patches *at time t* disperse to the other patch. Shoemaker et al. (2022) set the parameters in the environmental fluctuation functions *μ*_*x*_(*t*) and *μ*_*y*_(*t*) so as to generate variability on a short timescale in patch *x*, and conversely, on a long timescale in patch *y*. Further, they assumed that the growth rates of both species respond in the same way to the environmental fluctuation in patch *x* (i.e., *ε*_*x*,1_ = *ε*_*x*,2_) but respond oppositely in patch *y* (i.e., *ε*_*y*,1_ = *−ε*_*y*,2_).

As summarized in their Fig. 4, Shoemaker et al. (2022) showed that in the absence of dispersal, in patch *x*, where the short-timescale environmental driver affects each competitor in the same way, the two populations fluctuate synchronously on both short and long timescales (Fig. 2 a,j), a result that is also captured by the classic variance ratio. In contrast, in patch *y*, where the long-timescale environmental driver affects the competitors oppositely, compensatory dynamics are indicated on both timescales and by the classic ratio (Fig. 2 d,j).

**Figure 2.**
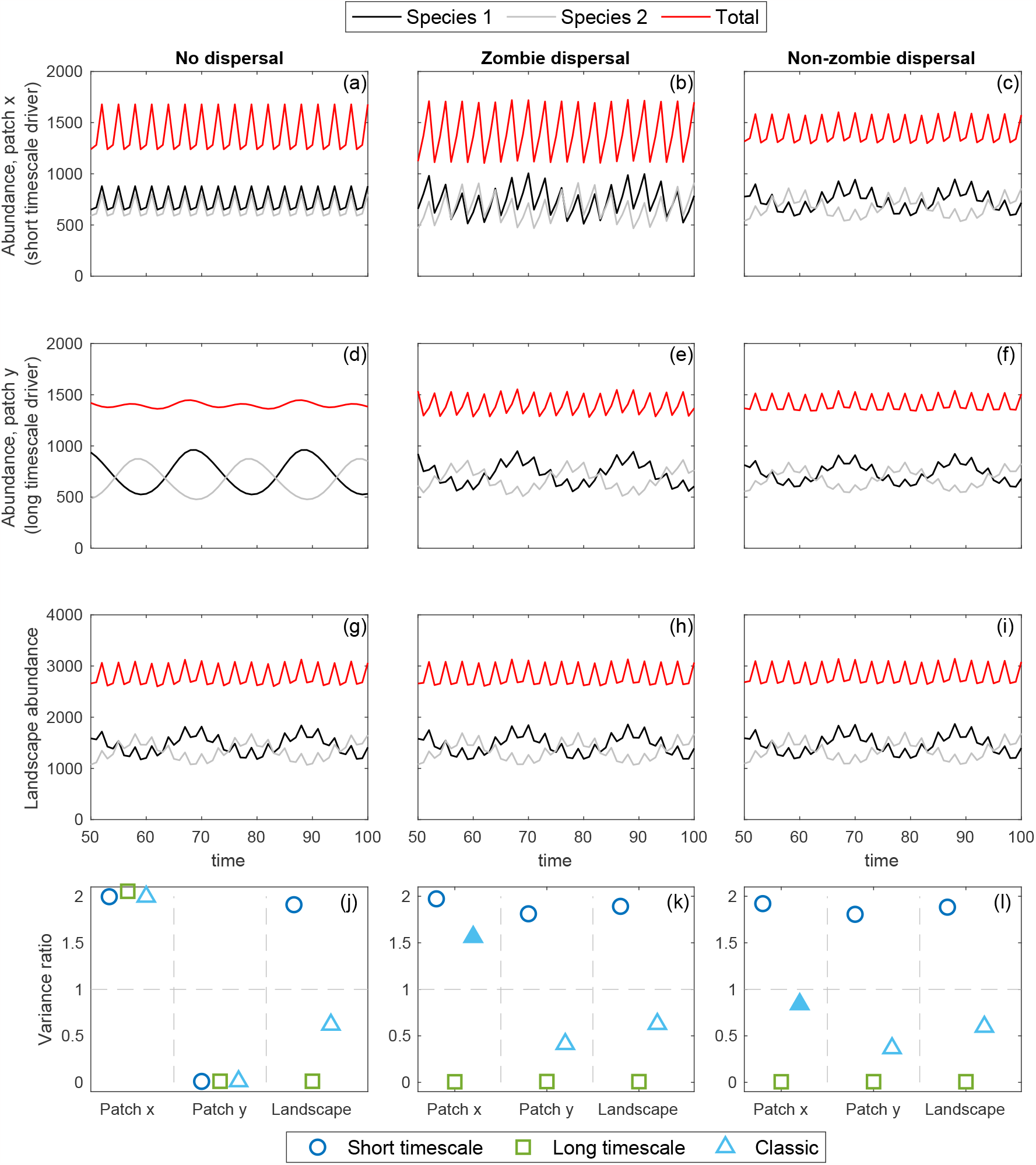
Species abundance in patch *x* (top row), patch *y* (second row), and aggregated across patches (third row), in models with no dispersal between patches (first column), with “zombie dispersal” (second column), or with “non-zombie dispersal” (third column). Timescale specific variance ratios at short (blue circles) and long (green squares) timescales, along with the classic variance ratio (cyan triangles) within patches and aggregated over both patches (landscape) are in the bottom row. Filled symbols indicate qualitatively different outcomes between zombie and non-zombie dispersal. Parameter values are: *a*_1_ = 0.5, *b*_1_ = 2*π/*3, *c*_1_ = 2, *a*_2_ = 1, *b*_2_ = *π/*10, *c*_2_ = 0, *r*_1_ = *r*_2_ = 0.5, *β*_1,2_ = *β*_2,1_ = 0.5, *K*_1_ = 1100, *K*_2_ = 1000, *ε*_*x*,1_ = *ε*_*x*,2_ = 0.5, *ε*_*y*,1_ = − *ε*_*y*,2_ = 0.1. In the “no dispersal” model *d* = 0; in the other models *d* = 0.4. Initial species abundances in each patch were *N*_*i,x*_ = *N*_*i,y*_ = *K*_*i*_. The parameter values and figure design parallel Fig. 4 of Shoemaker et al. (2022).

Dispersal, as described by model (7), changes the character of the dynamics in both patches (Fig. 2b,e): The signal of long-timescale compensation from patch *y* also appears in patch *x* and the signal of short-timescale synchrony in patch *x* also appears in patch *y* (Fig. 2k). The classic ratio in each patch reflects the effects of the environmental driver in that patch (synchronizing in patch *x* and compensating in patch *y*).

The question arises whether the results of Shoemaker et al. (2022) hold in a model that does not suffer from zombie dispersal. We can construct a “non-zombie” model by carefully accounting for the order of events. If dispersal is the last event before population census, the correct, non-zombie, model is

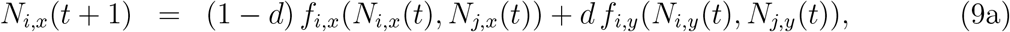

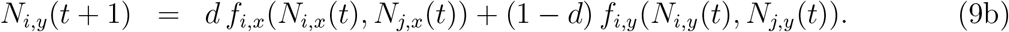

In the next section, we compare timescale specific variance ratios for the zombie and non-zombie dispersal models, and find that in many cases they lead to different conclusions about the effects of dispersal on synchrony.

## Simulation Results

We simulated both models (7,8) and (9,8) with the parameter values, initial conditions, and number of transient iterations as those in Shoemaker et al. (2022). We were able to duplicate Shoemaker et al.’s results with model (7,8) (Fig. 2, first two columns from left) showing that the classic variance ratio is representative of the dynamics of the focal patch (synchronous in patch *x* and compensatory in patch *y*) (Fig. 2k). With model (9,8), however, the result is different: the classic variance ratio indicates weak compensation in patch *x*, reflecting the significant difference in the dynamics there (compare Fig. 2b with Fig. 2c). The large amplitude oscillations in patch *y* (Fig. 2d) are now apparent in patch *x* (Fig. 2c) due to dispersal, as might be expected.

The different effects on synchronization generated by zombie vs. non-zombie dispersal are even more apparent under different ecological scenarios. In a high-reproduction scenario, zombie dispersal leads to (spurious) large fluctuations in population abundance in both patches (Fig. 2, b and e) and synchronizes the dynamics on both long and short timescales in patch *y* (Fig. 2, j and k). It has no apparent effect on the degree of synchronicity in patch *x*. In contrast, non-zombie dispersal dampens the fluctuations in patch *x* (Fig. 2, a and c) and reduces the differences in the population sizes of the two species in patch *y* (Fig. 2, d and f). The result is that non-zombie dispersal generates compensatory fluctuations on long timescales in both patches and over the landscape (Fig. 2l), the opposite of the effect of zombie dispersal.

The variance ratios (5) and (6) can also be used to investigate the ways dispersal affects cross-patch synchrony in the dynamics of the same species in the two different patches (e.g., the synchrony of *N*_1,*x*_(*t*) and *N*_1,*y*_(*t*)). For the parameters studied by Shoemaker et al. (2022), the zombie dispersal model synchronizes the spatial dynamics of both species on long time scales (Fig. 3: a and b, d and e, g and h), as might be expected. Zombie dispersal has only a very minor effect on cross patch synchrony as diagnosed by the classic ratio, and perhaps surprisingly generates compensatory cross patch dynamics on short timescales (Fig. 3 h). In contrast, non-zombie dispersal strongly synchronizes cross-patch dynamics for both species on both short and long timescales (Fig. 3 i). This difference between the effects of zombie versus non-zombie dispersal on intraspecific spatial synchrony is even stronger when population growth rates (Fig. 4 a-c) or dispersal rates (Fig. 4 d-e) are large.

**Figure 3.**
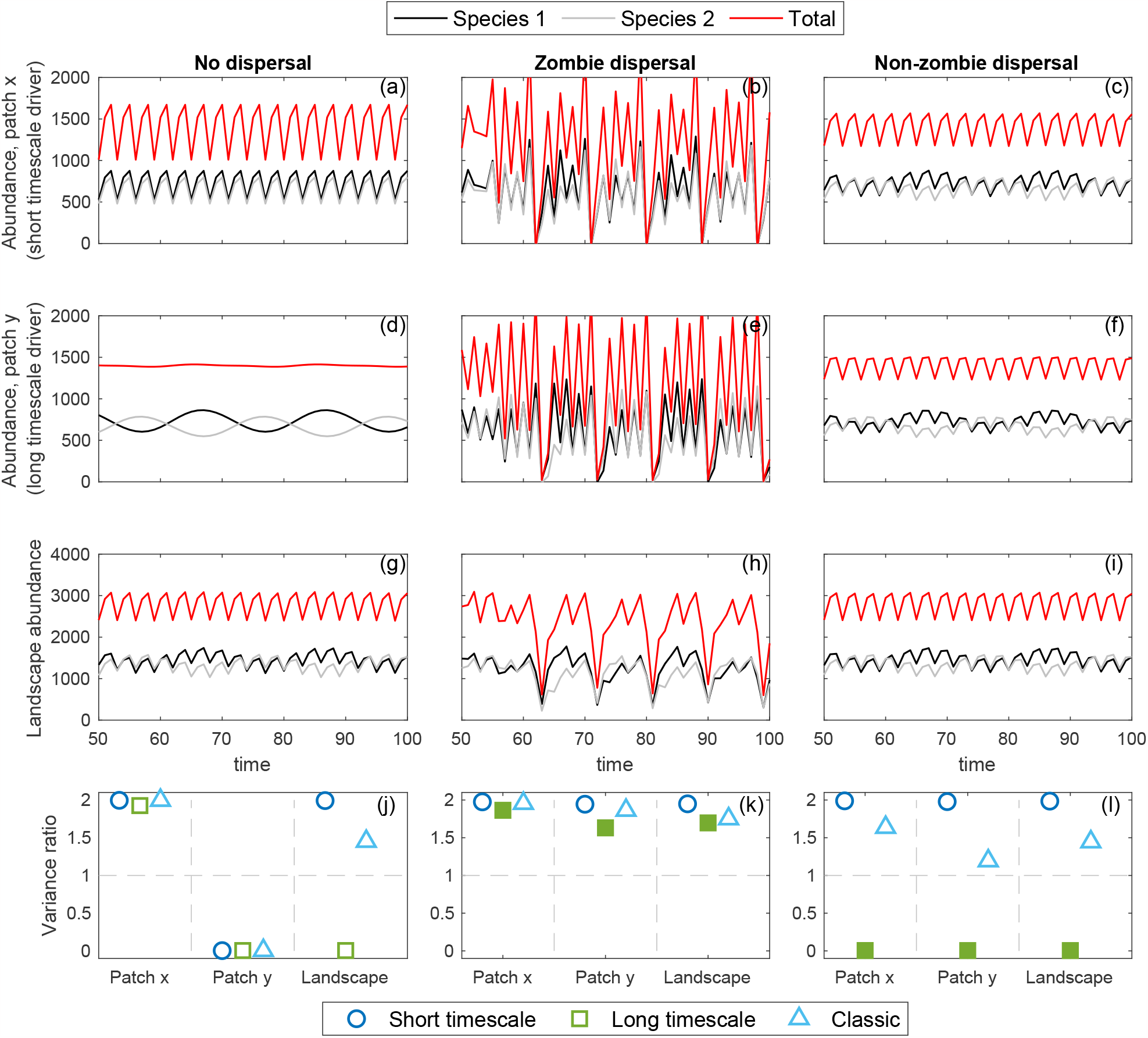
Species abundance in patch *x* (top row), patch *y* (second row), and aggregated across patches (third row), in models with no dispersal between patches (first column), with “zombie dispersal” (second column), or with “non-zombie dispersal” (third column). Timescale specific variance ratios at short (blue circles) and long (green squares) timescales, along with the classic variance ratio (cyan triangles) within patches and aggregated over both patches (landscape) are in the bottom row. Filled symbols indicate qualitatively different outcomes between zombie and non-zombie dispersal. Parameter values are as in Fig. 1, except *r*_1_ = *r*_2_ = 1.55.

**Figure 4.**
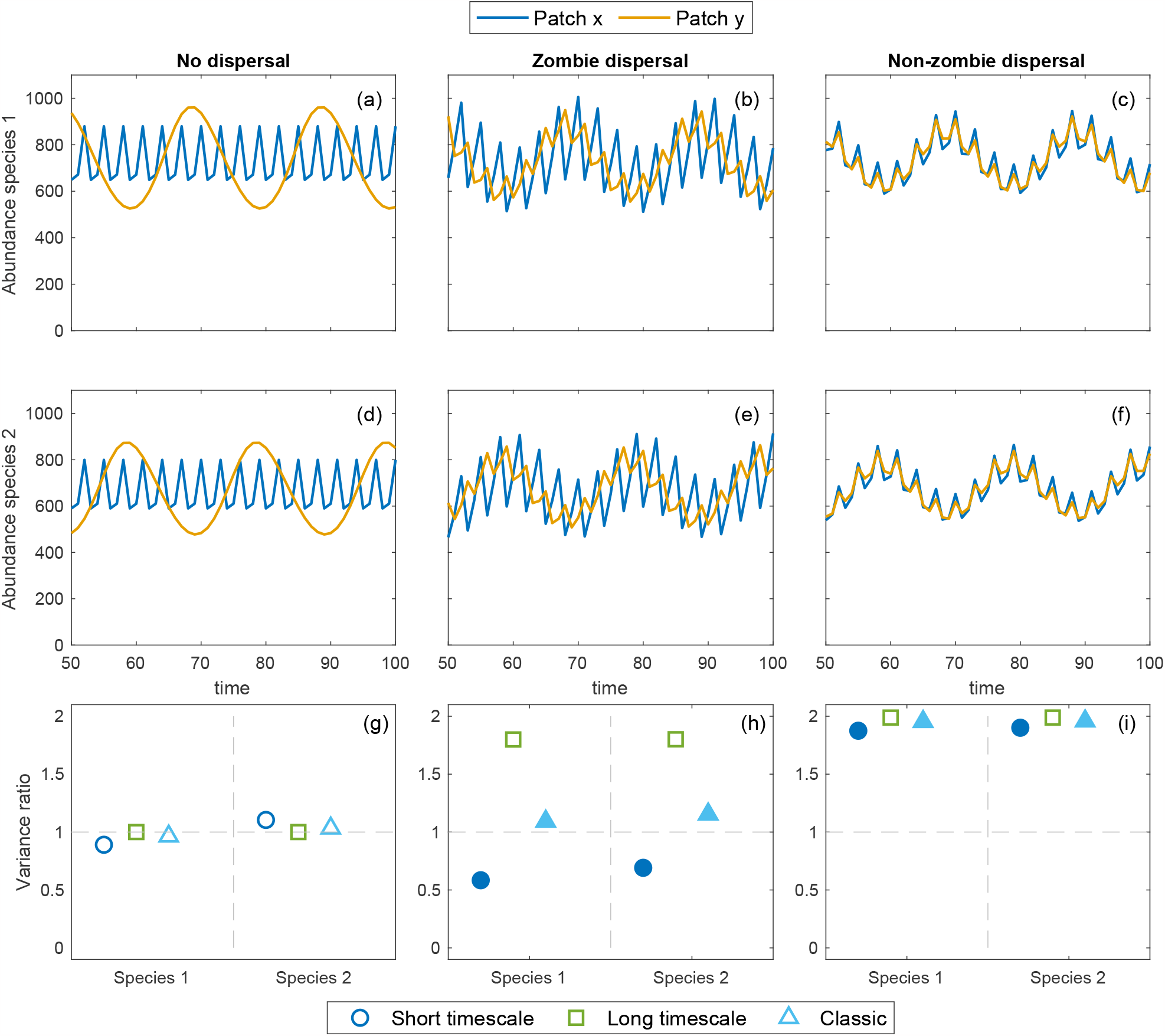
Population trajectories of species 1 (top row, a-c) and species 2 (second row, d-f), under the three dispersal models (no dispersal: a,d; zombie dispersal: b,e; non-zombie dispersal: c,f). Cross-patch timescale specific variance ratios (short timescale: blue circles; long timescales: green squares) and classic ratio (cyan triangles) for both species (bottom row, g-i). Filled symbols indicate qualitatively different outcomes between zombie and non-zombie dispersal. Parameter values are the same as in Fig. 1.

## Conclusion

We are not the first to draw attention to the zombie-dispersal problem. As noted by Lloyd (1995), Kot and Schaffer (1986) alerted readers that “discrete-time population models with linear coupling [a.k.a. zombie dispersal] in effect mix generations and, as a result, may yield counterintuitive results” and Jackson (1990) pointed out that the diffusive process must be kept separate from the reproductive process in order to “keep in touch with reality.” Hassell et al. (1995) warned that zombie dispersal can lead one to incorrect, counterintuitive results (e.g., that dispersal in a single-species model could be destabilizing), and emphasized “…how important it is that assumptions about mortality and dispersal are properly ordered in the organisms’ life cycle.” These warnings bear repeating, and generalizing: the careful ordering of all life history events is crucial for the formulation of logically-consistent non-linear discrete-time models. Further, different orderings can generate qualitatively different population (Neubert and Caswell, 2000) and metapopulation dynamics (Doebeli, 1995), evolutionary outcomes (Johst and Brandl, 1997), and management recommendations (Darwin and Williams, 1964; Bodine et al., 2012; da Silveira Costa and dos Anjos, 2019).

We note in passing that there is a growing literature on the analysis of mathematical models in which zombie dispersal is a *feature* rather than a problem. Indeed, mathematical ecologists have explored the effects of other zombie processes (see, e.g., Smith?, 2014), and have generated a sub-genera of mathematical zombie ecology and epidemiology. Of course, timing affects the effectiveness of control measures in discrete-time zombie models too (Ashlock et al., 2014)! The mathematical modeling of zombies has also been recognized as an effective way to engage students, public health professionals, and the general public (Lofgren et al., 2016). The subject just won’t die.

In our simulations, zombie dispersal artificially generates synchrony in the dynamics of the competing species within patches (Figs. 2 and 3), particularly on long timescales. In contrast, non-zombie dispersal is much more synchronizing than zombie dispersal when comparing the same species across patches (Fig. 4 and 5). Further exploration of this model may reveal what underlying configurations result in different scenarios of spatial and compositional variability (see Lamy et al., 2021). That the classic variance ratio fails to recover the synchrony signal of the patch-specific driver under non-zombie dispersal strengthens the argument of Shoemaker et al. (2022) for using a timescale-specific variance ratio in metacommunity settings.

**Figure 5.**
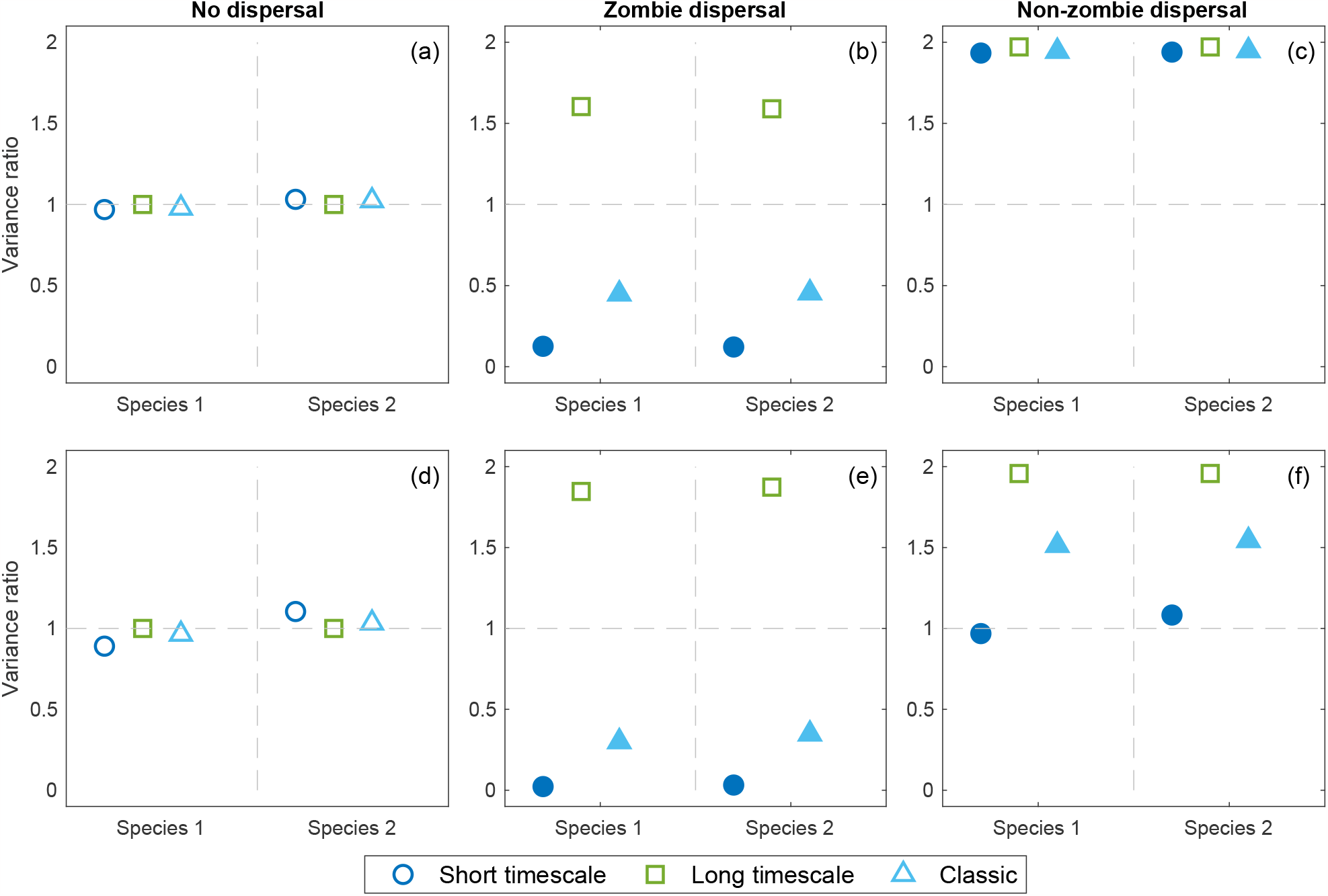
Cross-patch timescale specific variance ratios at short (blue circles) and long (green squares) timescales, along with the classic variance ratio (cyan triangles) for both species and three dispersal models. Parameter values are the same as in Fig. 1, except as follows: in panels (a)-(c) intrinsic growth rate is *r*_*i*_ = 1.55; in panels (d)-(f), the dispersal parameter is 0.85. Filled symbols indicate qualitatively different outcomes between zombie and non-zombie dispersal.

## Acknowledgements

The authors thank Heidi Sosik for discussions and suggestions that improved the final analysis and manuscript. This material is based upon work supported by the National Science Foundation (OCE-1655686) and by the Simons Foundation (561126).

## Footnotes

1 We have used the symbols *𝓊*_*x*_ and *𝓊*_*y*_ for reasons that will be made clear later.

2 Our choice of *𝓊* to represent population size in this model should now be clear; *𝓊* includes the “**u**ndead.”

3 Hastings (1993), Lloyd (1995), Kendall and Fox (1998), and many others have studied the dynamic of these equations when the local dynamics are logistic (i.e., *g*(*n*) = *rn*(1 − *n*)). Lloyd (1995) compares the dynamics of the two-patch logistic model with both zombie (linear) and non-zombie (nonlinear) dispersal.

